# Selective loss of Y chromosomes in lung adenocarcinoma modulates the tumor immune environment through cancer/testis antigens

**DOI:** 10.1101/2024.09.19.613876

**Authors:** Jonas Fischer, Katherine H. Shutta, Chen Chen, Viola Fanfani, Enakshi Saha, Panagiotis Mandros, Marouen Ben Guebila, Joanne Xiu, Jorge Nieva, Stephen Liu, Dipesh Uprety, David Spetzler, Camila M. Lopes-Ramos, Dawn L. DeMeo, John Quackenbush

## Abstract

There is increasing recognition that the allosomes, X and Y, play an important role in health and disease beyond the determination of biological sex. A loss of the Y chromosome (LOY) occurs in most solid tumors in males and is often associated with worse survival, suggesting that LOY may give tumor cells a growth or survival advantage. We here use an expression-based continuous measure of LOY that allows us to investigate LOY in lung adenocarcinoma (LUAD) using both bulk and single-cell expression data. We find evidence suggesting that LOY affects the tumor immune environment by altering cancer/testis antigen expression and consequently facilitating tumor immune evasion, also reflected in inferred gene regulatory networks. In immunotherapy data, we further show that LOY and changes in expression of particular cancer/testis antigens are associated with response to pembrolizumab treatment and outcome. This computational study provides new insights into the mechanisms behind LOY in LUAD and a powerful biomarker for predicting immunotherapy response in LUAD tumors in males.

## 1 Introduction

Beyond its role in male sex determination, the Y chromosome has been greatly understudied. Mounting evidence suggests that the Y chromosome is a driver of men’s health (Maan et al, 2017), and that loss of the Y chromosome (LOY) profoundly affects both health and disease (Wang and Sano, 2024). For example, LOY in blood has been shown to lead to cardiac fibrosis and increased heart failure mortality in mouse models (Sano et al, 2022), to be associated with higher risk and shorter survival in human cancers (Wright et al, 2017; Thompson et al, 2019; Forsberg et al, 2014), and with shorter lifespans in human males (Loftfield et al, 2018, 2019). Others have reported that LOY in the blood occurs naturally in aging men (Forsberg et al, 2014) and that LOY increases with smoking (Dumanski et al, 2015).

The functional consequences of LOY in blood have been reasonably well-studied. LOY in solid tumor tissue, however, has only recently been subject to deeper investigation. Early reports highlighted LOY in various cancer types, including non-small-cell lung cancer (Center et al, 1993), bladder cancer (Sauter et al, 1995), and papillary renal cell carcinoma (Brunelli et al, 2003), but the mechanisms driving LOY have remained elusive. Recently, investigations of large pan-tissue resources have found that LOY is evident in most solid tumors but not the corresponding healthy tissue, and that LOY can be associated with reduced cancer survival (Qi et al, 2023; Müller et al, 2023).

LOY has also been suggested as a potential driver event for cancer (Cáceres et al, 2020; Qi et al, 2023; Müller et al, 2023), but few studies have yet investigated this direct connection. Cáceres et al (2020) reported an extreme down-regulation of Y-chromosome genes that they hypothesize as a functional mediator between LOY and cancer but did not establish a direct connection. Both Qi et al (2023) and Müller et al (2023) investigated LOY in solid tumors using data from TCGA, where Qi et al (2023) provided the first systematic catalog of LOY in cancer, showing that LOY is a frequent somatic event in primary tumors. Considering both gene-expression-based “functional” LOY and full and partial LOY estimated through CNVs, they further showed that TP53 mutations are enriched in particular LOY-affected cancers and that LOY is a driver event for uveal melanoma. Müller et al (2023) demonstrated that a binary LOY indicator derived from CNVs is correlated with tumor mutational burden across cancers and provided further evidence that TP53 mutations are associated with LOY in some cancers, including LUAD. More recently, a functional study showed that LOY in mouse models of bladder cancer causes changes in the tumor immune microenvironment (TIME) and increased tumor growth (Abdel-Hafiz et al, 2023), providing a causal connection between LOY and tumor growth, possibly explaining the poor prognosis they found associated with LOY in human patients with bladder cancer. Overall, there is growing evidence that LOY is a driver event for cancer through its association with changes in the TIME. However, potential reasons behind how LOY achieves these alterations remain unclear.

In our study, we focus on lung cancer, which is estimated to be the most frequently diagnosed cancer and leading in cancer mortality, accounting for an estimated 1.8 million deaths in 2022 (Bray et al, 2024), with Adenocarcinoma (LUAD) as the most common lung cancer subtype.Although LOY has not been extensively studied in LUAD, it has been estimated that approximately 32% of LUAD tumors exhibit LOY; a result that is based on an assumed detection threshold. Here we present a computational analysis of LOY in individuals with and without LUAD (Figure 1) using bulk and single-cell sequencing from The Cancer Genome Atlas (TCGA), the Genotype-Tissue Expression project (GTEx), and the Human Tumor Atlas Network (HTAN). We suggest an effective expression-based metric that quantifies the LOY, allowing us to study LOY even when CNV or WGS data are not available, which is typical in clinical settings. Based on our metric, we identify a characteristic pattern of LOY in TCGA LUAD tumor tissue that reveals strong associations with immune cell composition in the tumor surroundings.We then provide evidence that LOY is selected for rather than being the result of a random chromosomal aberration and show that LOY does not occur in infiltrating immune cells. One of our key findings is that antigen presentation-related processes differ between cells with and without LOY, which provides a direct link between the LOY and TIME variability.

**Fig. 1.**
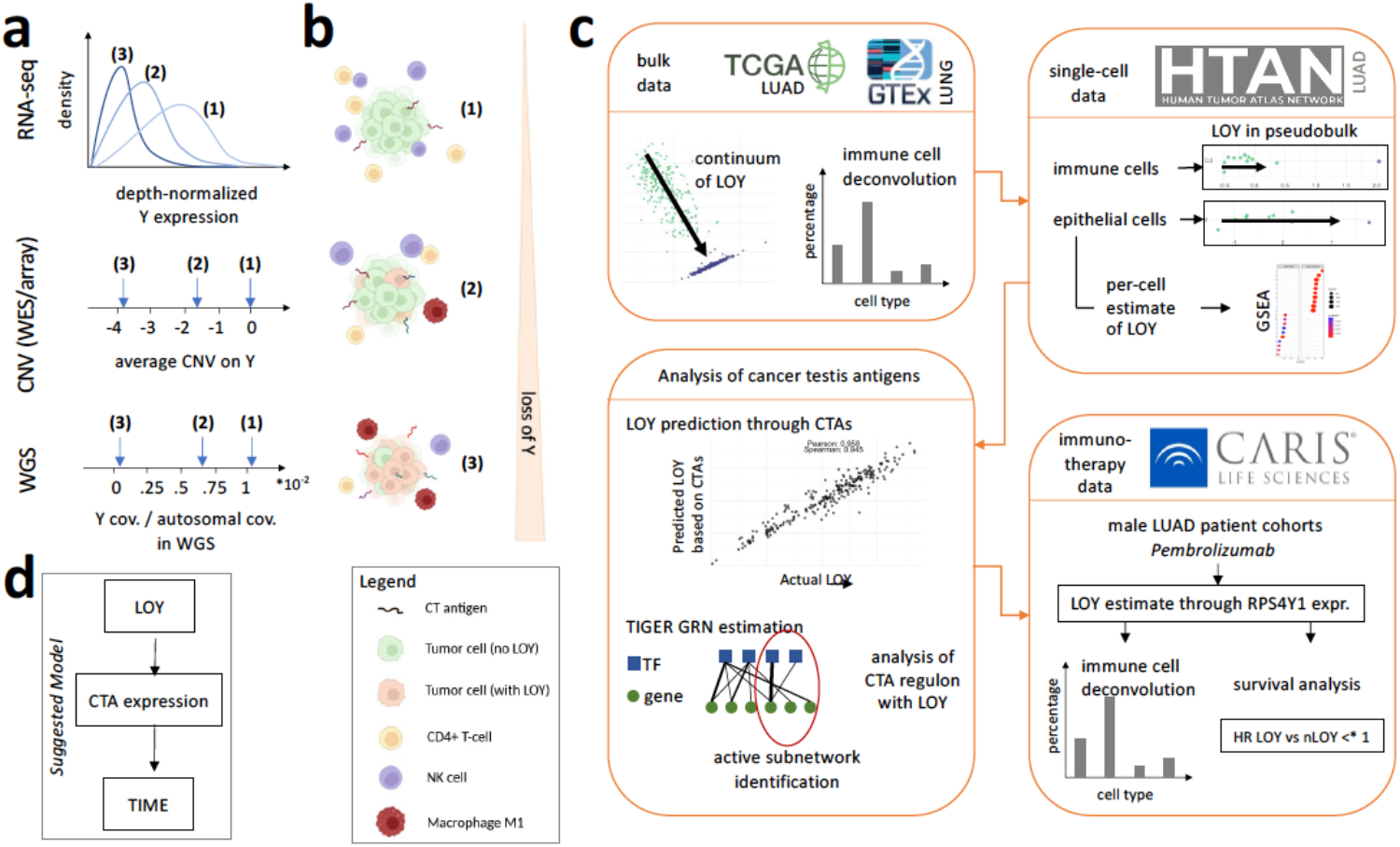
Study overview and key results. **a** Different *measures of LOY* including RNA-seq, copy number variant (CNV) estimates, and ratios of Y coverage to autosome coverage from whole genome sequence and how increasing ratios of LOY to non-LOY cells in the primary tumor would show in such measures (1: no LOY, 2: low LOY, 3: high LOY). **b** Changes in the tumor immune microenvironment (TIME) with increased LOY in the primary tumor. With increasing LOY we observed a change in TIME associated with alterations in the expression of some cancer/testis antigens (CTAs). The amount of NK, and CD4+ T-cells decreases with LOY, while M1 macrophage proportions consistently increase, steady proportions of immune cells are not shown. Partially created in BioRender BioRender.com/v90i141. **c** Overview of this study. Using lung adenocarcinoma as a model, we investigate *continuous* measures of LOY, demonstrate that this is driven by tumor and not immune cells, identify an association between LOY and cancer testis antigens and use gene regulatory network inference to analyze their expression, and then explore how immunotherapy response changes with LOY. **d** We suggest a mechanistic model of LOY modulating the TIME through CTAs.

We find that autosomal cancer/testis antigens (CTAs) are highly correlated with the degree of LOY and, using gene regulatory network modeling, demonstrate that differences in CTA regulation by transcription factors are linked to LOY. Cancer/testis antigens (CTAs) are proteins encoded by a group of autosomal genes that are primarily expressed in the testis, an immune-privileged site. They are, however, aberrantly expressed in many tumor types where they present as antigens to the immune system, initiating changes in the Tumor Immune Microenvironment (TIME) (Kortleve et al, 2022) and altering immune response. This study provides the first description of a process in which tumor cells exhibiting LOY may be selected for through a direct link between LOY, CTAs, and processes contributing to immune evasion.

The alterations in immunogenicity associated with LOY led us to speculate whether increased LOY, coupled with changes in CTA expression and associated changes in the tumor’s immune environment, could affect the efficacy of immunotherapy. We investigated the relationship between immunotherapy outcomes and differential expression of Y-genes and CTAs in male cohorts treated with pembrolizumab, a checkpoint inhibitor that targets the PD-1/PD-L1 pathway and one of the most commonly used immunooncology agents. We found that pembrolizumab treatment has a significantly better survival outcome in individuals with low *RPS4Y1* expression (the best single-gene indicator for LOY we found in TCGA LUAD cases). Furthermore, several CTAs for which we found significant regulatory changes with LOY also show significant association with immunotherapy survival outcomes, providing further evidence of CTA-mediated effects of LOY on the TIME. Our results suggest that LOY measures may serve as a biomarker for the directed selection of immunotherapeutic treatments for individual patients and that CTA-targeting immunotherapy may complement these treatments to improve overall treatment outcomes.

## 2 Methods

### 2.1 Data Sources

#### Bulk RNA-seq data

We obtained publicly available bulk RNA-sequencing data from the Genotype-Tissue Expression (GTEx) project and The Cancer Genome Atlas (TCGA). In both cases, data were uniformly reprocessed using `recount3` (Wilks et al, 2021), version 1.8.0. Raw counts were normalized by conversion to transcripts per million (TPM); a pseudocount was added, and data were log (base *e*) transformed as log(*TPM* + 1). TCGA data were separated into adjacent normal and primary tumor tissue. Duplicates were removed, keeping only the sample with the greatest sequencing depth. RNA-seq based sample purity estimates were obtained from Aran et al. (Aran et al, 2015).

#### Whole-genome sequencing data

Whole-genome sequencing (WGS) alignment files of N=278 male LUAD patients in TCGA were accessed through dbGaP (list of IDs given in Supplementary Table 1). Coverage of Y and autosomal genes was computed using samtools (Li et al, 2009), version 1.18. To facilitate the comparison of WGS-based LOY estimates with gene expression-based LOY estimates, we considered only high-coverage WGS samples that had at least 50% of the maximum observed coverage across samples (N=57).

#### Calculating WGS-based loss of Y in GTEx and TCGA

LOY estimates from WGS are computed through the number of reads covering Y divided by the number of reads covering autosomes, which gives a unit-less relative measure of the loss of chromosome Y. Assuming that reads are distributed uniformly at random over all chromosomes, potential individual biases in read depth were thus normalized to provide a standardized measure of LOY.

#### Calculating CNV-based LOY in TCGA

To obtain copy number variation (CNV)-based LOY estimates, we used the `TCGAbiolinks` package (Colaprico et al, 2016), version 2.25.3, to retrieve copy number segment information for CNVs. LOY scores are based on the mean CNV value across segments on the Y chromosome.

#### Single-cell expression data

Single-cell RNA-seq data for lung adenocarcinoma from 8 male and 16 female patients were obtained from the Human Tumor Atlas Network (HTAN) (Chan et al, 2021) through CELLxGENE (Abdulla et al, 2023) on 02/24/2023, which had already been processed using `Seurat` (Stuart et al, 2019). Data were split into epithelial and immune cells according to the provded cell-type annotation and filtered for lung tumor tissue. To conduct subject-level analysis, single-cell data was pseudobulked by summing read counts across cells of the same individual, for each cell normalizing the count by the total number of overall reads in this cell (read depth). For subject-level normalization, we divided by the number of cells for this subject. As validation data, single-cell LUAD expression data from Kim et al (2020) was downloaded through GEO (GSE131907), and the same procedure of filtering for lung tissue, partitioning cells into epithelial and immune groups, and pseudobulking was applied.

### 2.2 Gene expression-based LOY estimation

#### The pc-LOY estimate

To create a measure for LOY based only on bulk gene expression, we first compute the principal components (PCs) of the Y-chromosome RNA-seq data within each dataset. The PCs are computed on data containing both male and female samples; female samples serve as an anchor point for gene expression in individuals without a Y chromosome. We excluded Y genes in pseudoautosomal regions (PAR1 and PAR2) (Ross et al, 2005).

In TCGA LUAD, projecting the samples onto the first three PCs produces a clear separation of the data into two clusters corresponding to male and female samples (Supplementary Figure 1a). These PCs account for 93.5% of the variance in the data (Supplementary Figure 1b).

To obtain the LOY score, in brief, we use the female individuals as anchor points– showing virtually zero Y expression–and define a direction from those towards the male individuals in this Y expression space. By mapping each male to this direction vector, we have an estimate of how close a male individual is to the female anchor points, in other words, how much LOY that particular individual has undergone. More formally, we take the vector connecting the female centroid *µ*_*F*_ ∈ ℝ^3^ to the male centroid *µ*_*M*_ ∈ ℝ^3^. We map each male individual’s PC coordinates *p*_*i*_ ∈ ℝ^3^ to a univariate measure *q*_*i*_ ∈ ℝ by orthogonal projection onto this vector (Figure 2b), which is given by the inner product,

**Fig. 2.**
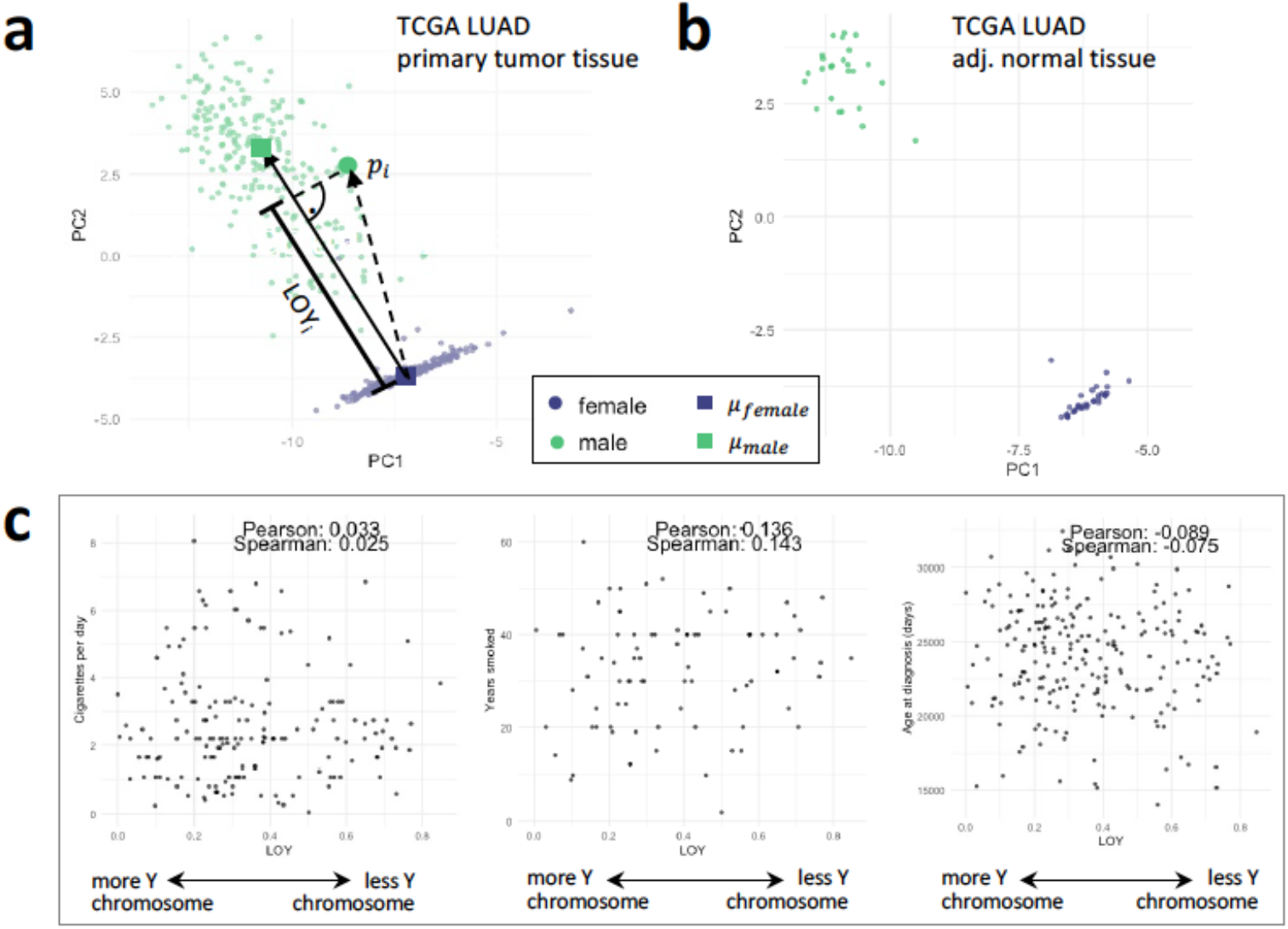
Loss of Y in LUAD. **a** First two principal components of Y genes in TCGA primary tumor tissue of LUAD patients, providing evidence for LOY based on gene expression levels. Overlaid, we visualize our approach to estimate LOY of primary tumors from Y gene expressions based on the first three principal components of patient i (third principal component not shown here for ease of understanding). **b** First two principal components of Y genes in TCGA adjacent normal tissue of LUAD patients showing no continuity of LOY compared to tumor tissue. **c** Covariates with known association in blood LOY show no association in tumor LOY. From left to right: Self-reported cigarettes per day, self-reported years smoked, and reported age at diagnosis against LOY estimate in TCGA LUAD.

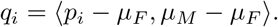

We then transform these PC-based values to a [0, 1] measure of LOY (“pc-LOY”) by scaling to the maximum observed LOY value,

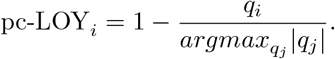

#### Association of pc-LOY with covariates

We analyzed the association of pc-LOY with covariates available as annotated metadata from `recount3` (Wilks et al, 2021), version 1.8.0. Specifically, we considered age at diagnosis, batch effect as given by the annotated CGC case batch number, smoking exposure as indicated by the number of years smoked, and smoking exposure as indicated by cigarettes per day. We further considered sample purity estimated through ESTIMATE (Aran et al, 2016). We used the pre-computed values for TCGA by Aran et al (2016).

### 2.3 Analysis of LOY

#### Predicting LOY with cancer/testis antigen expression

We retrieved N=266 annotated cancer/testis antigens (CTAs) from CTdatabase (Almeida et al, 2009). Using the autosomal CTAs, we fitted a multivariate least-squares model to predict LOY for male samples from CTA expression levels.

#### Investigating regulation in the promoter region of CTAs

Motif matches for binding sites of the two Y chromosome transcription factors encoded by *SRY* and *HSFY2* and the androgen receptor in the promoter regions of CTAs were extracted from the human motif prior in the GRAND database v1.5.0 (Ben Guebila et al, 2021), which were derived from motif scans using FIMO (Grant et al, 2011) against gene promoter regions defined as ± 1kb around the transcription start sites annotated in human reference genome hg38. They define a motif for each TF using position weight matrices from the CIS-BP 1.94d database (Weirauch et al, 2014). The motif call was performed using an adjusted p-value cut-off of 10^−4^.

#### Predicting LOY through deconvoluted cell quantities

We predicted the pc-LOY as the response, taking all cell type quantities estimated through quanTIseq v3.16 (Finotello et al, 2019) as features in a multivariate least squares regression, with results presented in Supplementary Figure 1c.

#### Adjudicating LOY in single-cell data

To adjudicate LOY on a single-cell level from scRNA-seq, we define cells with less than 0.04% of total reads coming from the Y chromosome to be affected by LOY. All other cells were considered as non-LOY. This threshold was determined based on a dip in the coverage distribution (see Supplementary Figure 1d).

#### GSEA in single-cell LOY data

We conducted Gene Set Enrichment Analysis based on the log-fold-change between the mean observed gene expression in LOY and non-LOY cells. We use `clusterProfiler` (Yu et al, 2012; Korotkevich et al, 2021), version 4.10.0, with a minimum gene set size set to five and report pathways as enriched based on a p-value cutoff of 0.05 for Benjamini-Hochberg corrected p-values.

### 2.4 Gene regulatory networks

#### Estimating sample-specific gene regulatory networks

To estimate sample-specific gene regulatory networks for the TCGA LUAD data, we used TIGER (Chen and Padi, 2024) from NetZooR 1.5.4 (Ben Guebila et al, 2023), a Bayesian matrix factorization method for inferring regulatory networks that leverages prior knowledge of transcription factor (TF) binding to decompose a gene expression matrix into the product of an overall gene regulatory network and sample-specific TF activities. More specifically, TIGER decomposes a gene expression matrix *G* ∈ ℝ^*m×n*^ of *m* genes and *n* samples into *G* = *WZ*, where *W* ∈ ℝ^*m×t*^ is a gene regulatory matrix for the *t* TFs, specifying the regulatory effect of each TF on each gene, and 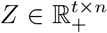 specifies the “availability” of each TF in each sample.

TIGER incorporates prior knowledge about the binding of TFs to gene promoters. Here, we consider signed TF-gene interactions with high confidence levels according to the DoRothEA database v1.12.0 (Garcia-Alonso et al, 2019), excluding TFs with support from only one computational resource at the DoRothEA level E, resulting in N=735 TFs in the prior. We considered the 16 856 protein-coding genes in the input expression matrix that were expressed in more than 25% of the samples. Using these genes and the 712 TFs overlapping with the gene set, including X and Y chromosomal TFs and genes, we applied TIGER to the set of N=193 males in the TCGA LUAD data to estimate the regulatory network *W* and individual TF activities *Z*.

To study individual-specific regulatory effects of TF *i* on gene *j* in a given sample *k*, we compute

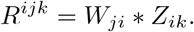

As an aggregate measure of the regulation explained by all TFs on gene *j* in a given sample *k*, we compute

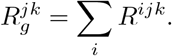

For assessment of the relationship between LOY and changes in gene regulation, we calculated the correlation between pc-LOY and 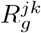 for each gene *j* across all samples *k*. Based on the 1000 genes exhibiting the regulatory changes most strongly correlated (in absolute value) with pc-LOY, we carried out a gene set enrichment analysis in KEGG pathways. The set of all 16856 genes used in the gene regulatory network was used as a background set. We report significantly over-represented pathways based on a p-value cutoff of 0.05 for FDR-corrected p-values.

We also analyzed individual CTAs and immune-related genes using gene-level regulatory scores *R*_*g*_. Here, we computed the Pearson correlation coefficient along with its t-test statistics between each 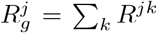, for CTA and immune-related genes *j*, summing over all samples *k*, and the pc-LOY. FDR multiple test correction was applied, and corrected p-values *<* .05 were reported as significant.

#### Active subnetwork detection

Active subnetwork detection uses graph search methods to identify subnetworks enriched for nodes associated with a particular outcome (Nguyen et al, 2019). To identify active subnetworks enriched for association with LOY in the TIGER LUAD GRN, we used a heuristic variation of the integer linear programming algorithm developed by Dittrich et al. (Dittrich et al, 2008) as implemented in the `runFastHeinz` function of the BioNet R package v1.49.1 (Beisser et al, 2010). This algorithm searches subgraphs in a node-weighted graph by maximizing a subgraph score based on the sum of the node weights; in this scheme, positive node weights are desirable. Following the approach of Dittrich et al. (Dittrich et al, 2008), we used p-values previously calculated for the association of each node in the TIGER network with the LOY trajectory. We fit a beta-uniform mixture (BUM) model to the distribution using the `fitBumModel` function and converted p-values into a score using the `scoreNodes` function, which assigns higher scores to lower p-values and viceversa. By design, the BUM model controls the FDR at a user-specified rate such that scores are positive for nodes below the FDR threshold and negative for nodes above it (Dittrich et al, 2008).

### 2.5 Immunotherapy data

Real-world clinical data were obtained from insurance claims, which encompass detailed records of health services, including prescribed medications, procedures performed, and established diagnoses. Time-on-treatment (TOT) was determined as the interval from the initiation to the conclusion of treatment. Kaplan-Meier survival estimates were generated for cohorts defined by molecular characteristics. Hazard ratios (HR) were computed using the Cox proportional hazards model, and significant differences in survival times were assessed with the log-rank test, where *P <* 0.05 was considered significant.

For RNA expression measurement, FFPE specimens underwent pathology review to measure percent tumor content and tumor size; a minimum of 10% of tumor content in the area for microdissection was required to enable enrichment and extraction of tumor-specific RNA. The Qiagen RNA FFPE tissue extraction kit was used for extraction, and the RNA quality and quantity were determined using the Agilent TapeStation. Biotinylated RNA baits were hybridized to the synthesized and purified cDNA targets, and the bait-target complexes were amplified in a post-capture PCR reaction. The Illumina NovaSeq 6500 was used to sequence the whole transcriptome from patients yielding an average of 60M reads. Raw data was demultiplexed by Illumina Dragen BioIT accelerator, trimmed, counted, PCR-duplicates removed, and reads were aligned to human reference genome hg19 using STAR aligner (Dobin et al, 2013). Transcripts per million (TPM) values were generated using the Salmon expression pipeline (Patro et al, 2017). RNA deconvolution was performed using quanTIseq method to characterize the tumor microenvironment (Finotello et al, 2019).

## 3 Results

### 3.1 A continuum of loss of Y in lung adenocarcinoma

Although microdeletions of Y-chromosomal segments are known, we considered LOY within a single tumor cell as a binary quantity reflecting whether or not, in genetically male individuals (those carrying both X and Y chromosomes), the Y chromosome is absent. In contrast, LOY in a collection of cells (such as a tumor biopsy) is a continuous measure reflecting the fraction of cells that have lost the Y chromosome. We defined the amount of LOY based on a linear mapping of principal components of Y gene expression in a given dataset (“pc-LOY”; Figure 2a; Methods). pc-LOY reflects the continuous nature of LOY in tumors and is general enough to apply to a wide variety of data sources, as it is solely based on bulk gene expression rather than CNV or WGS, often used in the literature.

In TCGA-LUAD, pc-LOY correlates well with Y-chromosome copy number variations (Pearson *ρ* = 0.74, *p <* 2.2 ∗ 10^−16^; Supplementary Figure 2a) and with estimates of chromosomal loss based on whole genome sequencing (WGS) as described in Methods (Pearson *ρ* = 0.78, *p* = 1.491 ∗ 10^−12^; Supplementary Figure 2b). We note that pc-LOY is a data-driven measurement that is specific to each dataset and is reestimated for every new cohort. Examining pc-LOY in non-tumor tissue reveals no evidence of LOY in either TCGA LUAD adjacent “normal” lung tissue samples or GTEx lung samples (Figure 2b, Supplementary Figure 2c).

#### LOY is a systematic loss in tumor cells

As LOY could be a random aberration due to the small size of the Y chromosome or its relatively sparse gene landscape. We thus compared it to the loss of chromosome 21, which is comparably small and gene-poor (compare Supplementary Figure 3a,b). In tumor tissue, we found no evidence of an association between chromosome 21 loss (defined as read coverage relative to genome-wide coverage) and LOY (Pearson *p* − *value <* 0.05), providing further evidence that LOY is a systematic loss compared to autosomal genes.

To rule out potential biases in the pc-LOY signal, we further tested four potential drivers and confounders of LOY in TCGA LUAD tumors—age, smoking history (cigarettes per day and years of smoking), sample purity estimates (Aran et al, 2015), and sample batches (Figure 2c, Supplementary Figure 3c,d)—and found no significant association between these variables and pc-LOY (Pearson correlation based t-test, p-value cutoff 0.05; Supplementary Table 2).

### 3.2 LOY in tumor cells alters immune cell interactions

Although bulk RNA-seq data have been instrumental in exploring the molecular mechanisms driving cancer, it does not capture the complex interplay between tumor and stromal cells. In males, LOY has been reported to be an age-associated phenomenon in white blood cells (Sano et al, 2022; Brown and Machiela, 2020; Cáceres et al, 2020; Qi et al, 2023; Müller et al, 2023), leading us to question whether the tumor-associated LOY we observed was driven by infiltrating immune cells. We analyzed single-cell RNA-seq data in LUAD tumors from the Human Tumor Atlas Network (HTAN) (Chan et al, 2021) separating the single-cell data into epithelial and immune cell groups based on the author-provided cell labeling. We constructed tumor and white blood cell pseudobulk datasets by pooling the respective scRNA-seq data and performed pc-LOY analysis on each separately. We found a LOY trajectory in epithelial cells, but not in immune cells within the tumor sample, indicating that LOY is tumor cell specific (Figure 3a); these results were replicated in LUAD single-cell data from Kim et al (2020) (Supplementary Figure 4). The evidence that LOY is present only in tumor cells – not in healthy GTEx samples, adjacent normal TCGA-LUAD tissue, or immune cells within the tumor microenvironment – suggests that changes in the microenvironment may reflect selective pressure associated with LOY in tumor cells.

**Fig. 3.**
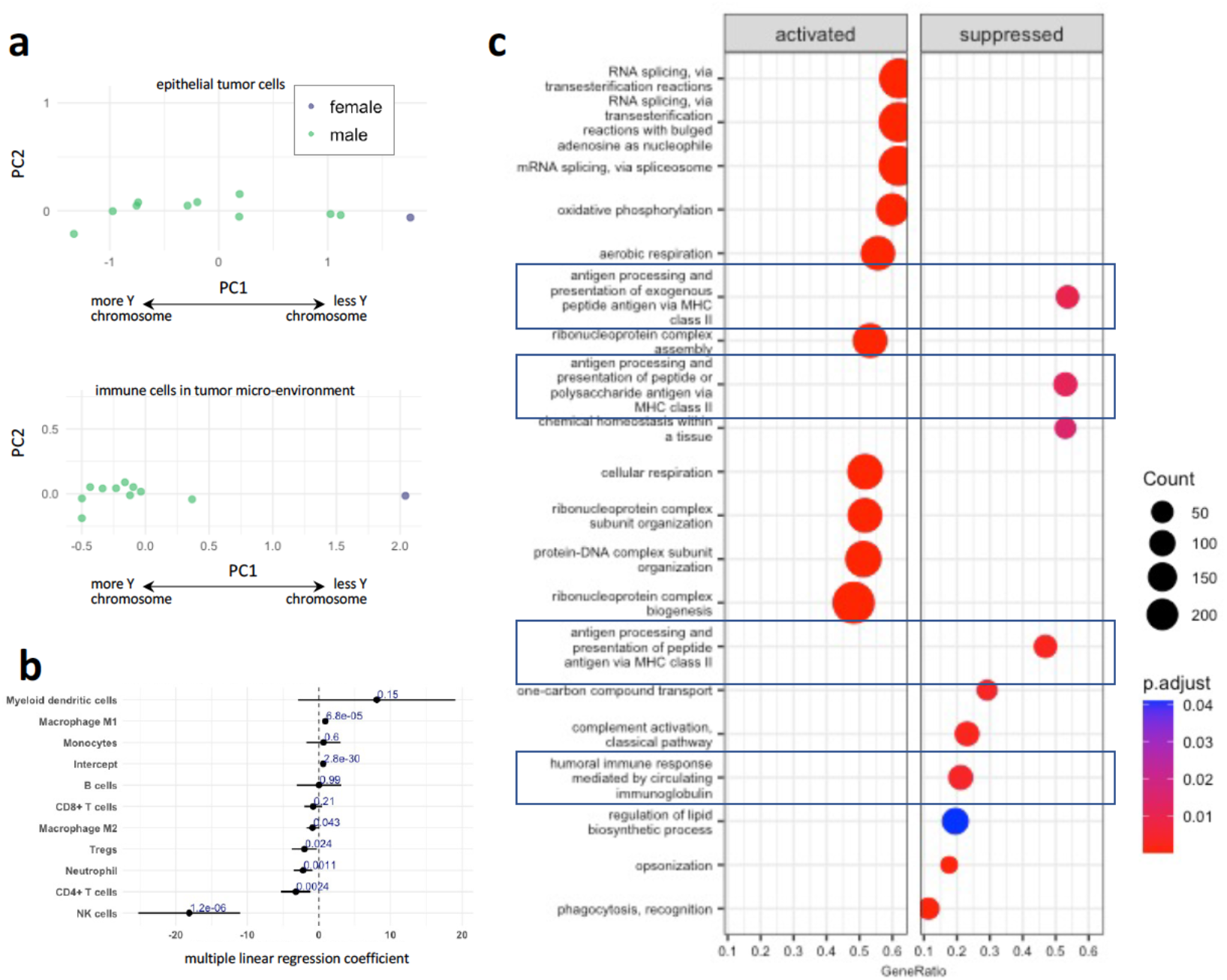
LOY and the immune system. **a** First two principal components on Y genes of pseudobulked gene expression of annotated epithelial cells and annotated immune cells for all samples in LUAD single-cell data. Female individuals collapse to a single point corresponding to zero expression. **b** Forest plot for coefficients of Quantiseq immune cell type fractions in least squares LOY fit. Visualized are coefficient estimates with standard error and p-value. **c** HTAN LUAD gene set enrichment analysis of GO terms based on log-fold change of gene expression between cells that lost Y and those that did not. Reported are the top 20 enriched terms with BH corrected *p <* 0.05. We see an enrichment of antigen-related pathways towards LOY-affected cells.

#### A selection for immune cell composition in the tumor

Abdel-Hafiz and colleagues reported that in mouse models of bladder cancer, LOY causes a different immune cell composition in the tumor microenvironment (Abdel-Hafiz et al, 2023). This led us to speculate that tumor immune invasion might select for LUAD cells that have lost their Y chromosome. Using TCGA LUAD data, we estimated the fractions of different immune cell types through computational deconvolution using quanTIseq (Finotello et al, 2019; Sturm et al, 2020). We examined the distribution of immune cell types with pc-LOY and found that immune cell make-up was predictive of LOY (least squares fit from multivariate regression of pc-LOY against all quanTIseq immune cell types, adj. *R*^2^ = .14, *p* = 3.143 ∗ 10^−6^). Significant predictors of LOY (univariate *p <* 0.01) in this model include M1 macrophages increasing in quantity with LOY, and NK cells, Neutrophils, and CD4+ T-cells decreasing in quantity with LOY (see Figure 3b). Both NK cells and CD4+ T-cells are of particular importance for tumor control and tumor immune response through mediation of cytotoxicity (Wu et al, 2020), with CD4+ T-cells explicitly recognizing antigens presented through HLA-II (Jhunjhunwala et al, 2021). In contrast to these significant predictiors, for non-significant predictors we usually found estimates close to zero for most of the samples. In the HTAN LUAD single-cell data, we found a similar trend, with the percentage (as a fraction of the total immune cell population) of both NK cells and T-cells decreasing with LOY. We used the available annotation from the original study; annotation for macrophages was not available within the HTAN LUAD data.

#### LOY-associated immune response on a process level

Abdel-Hafiz et al. also reported that changes in bladder cancer TIME may be induced through cellular programs associated with LOY. We used HTAN LUAD single-cell data to compare gene expression in tumor cells with and without LOY. We identified significant enrichment of GO biological processes related to antigen presentation in tumor cells with LOY (Benjamini-Hochberg (FDR)-corrected enrichment *p <* 0.05, Figure 3c). This supports the hypothesis that LOY may indeed induce changes in tumor cells that alter their interactions with immune cells.

#### Cancer/Testis Antigens as the mechanistic mediator

Immune cells recognize other cells by identifying antigens on their surfaces. Cancer/testis antigens (CTAs) (Fratta et al, 2011) are a unique group of antigens expressed only in the human germ cells of the testis (and occasionally in the placenta and ovary, all “immunoprivileged” sites), not in other normal tissues. However, they are also aberrantly expressed in various types of cancer (Maxfield et al, 2015; Jhunjhunwala et al, 2021). Several CTAs have been shown to have direct gene regulatory effects in cancer, such as *PAGE4* through modulation of the MAPK pathway (Lv et al, 2019), *CEP55* by regulating FOXO1 signaling (Zhang et al, 2022), and being linked with effects on cancer, with *FBXO39* knockdowns slowing down tumor growth in squamous cell tumors (Yang et al, 2022). *ADAM29* and its mutations significantly influence the proliferation, migration, and invasion of breast cancer cells (Zhao et al, 2016), and *MAGEC3* promotes epithelial-mesenchymal transition in esophageal squamous cell carcinoma (Wu et al, 2021). CTAs have also been shown to have an explicit T-cell response in testicular cancer (Pearce et al, 2017) and have been identified as a potential key component in the mechanisms of cancer immune evasion (Kortleve et al, 2022).

We investigated whether changes in CTA expression might explain the TIME changes we observed and found a significant correlation between autosomal CTA expression and LOY (least squares fit adj. *R*^2^ = 0.43, *p* = 0.0118). We then tested for a possible regulatory relationship between LOY and CTAs. Searching CTA promoter regions for binding sites of the Y chromosome-based transcription factors *SRY* and *HSFY2* and the Androgen Receptor (*AR*), which is regulated by the Y-encoded *KDM5D*, we find a gene regulatory mechanism of LOY affecting CTA expression. A complete list of CTA promoters containing binding sites for *SRY* and *HSFY2* can be found in Supplementary Table 3. We thus have a possible mechanistic link of LOY affecting the presence of specific antigens, which in turn affect tumors via TIME as well as gene regulation.

#### Gene regulatory network models of LOY support the evidence for TIME involvement

For a more principled approach to exploring the regulatory changes with LOY, we computationally inferred gene regulatory networks using TIGER. TIGER uses Bayesian matrix factorization to simultaneously infer transcription factor (TF) regulomes and TF activities from RNA-seq data (Chen and Padi, 2024). We computed TIGER gene regulatory networks for male TCGA-LUAD subjects (see Methods 2.4) and selected the 1000 genes with the greatest changes in expression that were explained by LOY-associated TF regulatory disruption. Gene set enrichment analysis identified significant (*p <* 0.05, FDR-corrected) enrichment of processes related to cell cycle and repair, processes reflective of tumorigenesis. Considering all autosomal CTA genes in the TIGER model, we find 29 of 266 for which expression variation by TF regulation is significantly correlated with LOY (t-test on Pearson correlation coefficient; FDR-corrected *p <* 0.05, Figure 4a). Among these is *XAGE1B*, which is among the most consistently expressed CTAs in the studied data and has been reported to trigger a humoral immune response in non-small cell lung cancer patients (Nakagawa et al, 2005), linking LOY and CTA expression changes in the TIME.

**Fig. 4.**
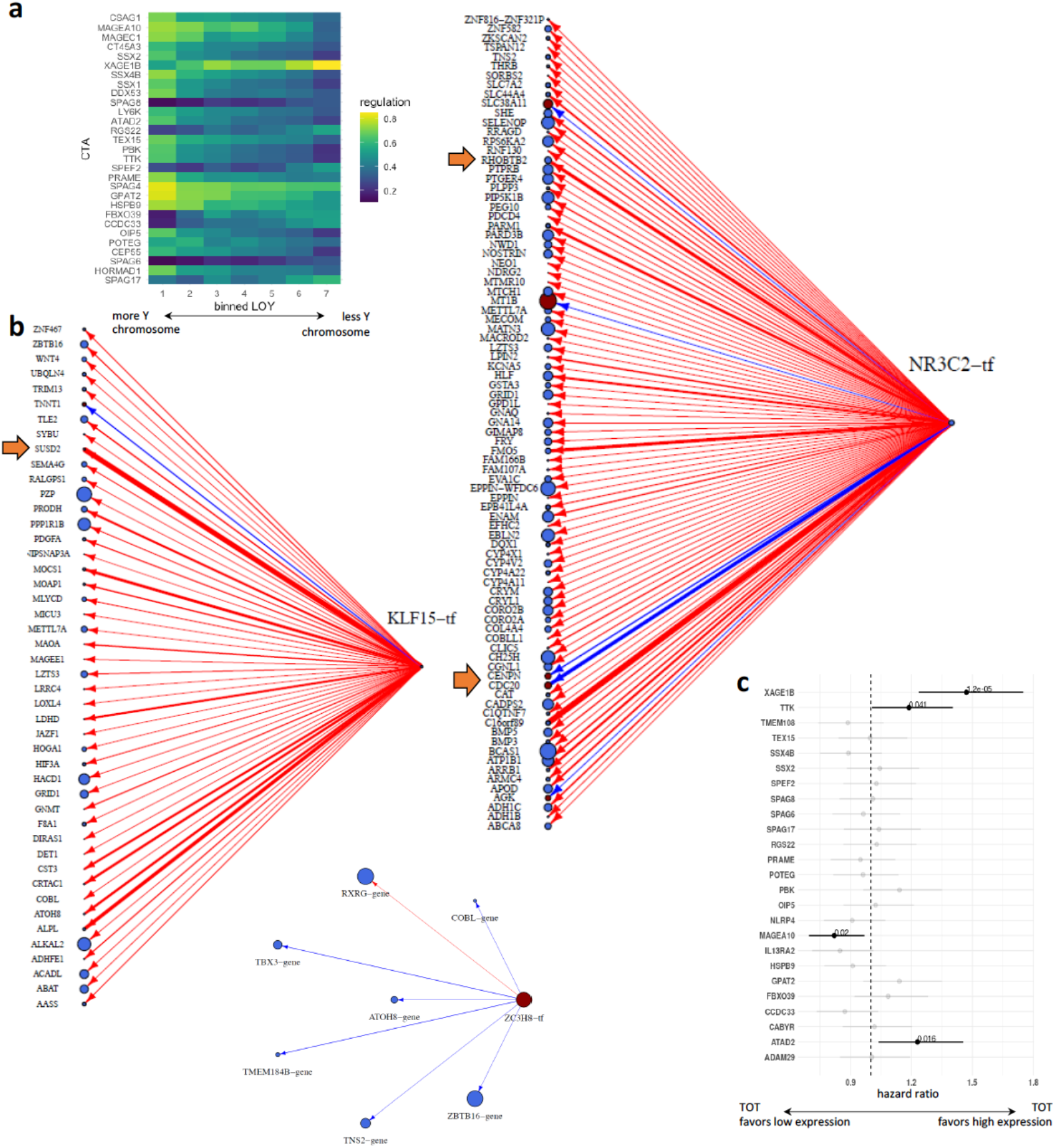
LOY and the immune system. **a** Regulatory effect of Cancer/Testis Antigens (CTAs) with strength of LOY within a LUAD gene regulatory TIGER network. To account for individual-specific noise, we construct seven equi-width bins across observed male pc-LOY, averaging networks across individuals within one bin. Visualized are CTAs that show significant correlation with LOY after BH correction. **b** Active subnetworks of TIGER network for three key TFs. Node size is proportional to the association of gene/TF with LOY, coloring is based on the type of association (blue: negative, red: positive). Edge line-width is proportional to weight in GRN, coloring is based on the regulatory effect (blue: negative, red: positive) **c** Hazard ratio (y-axis) when comparing male LUAD patients treated with pembro split into two groups based on median expression value of CTA (x-axis). Depicted are HR values along with confidence intervals and p-values in bold color; all insignificant observed HRs are transparent for clarity of the results.

To study the relationship between LOY and immune-related gene regulation in depth, we analyse the regulation of 31 known immune-related genes, including drivers of the immune response, pro-inflammatory cytokines, genes encoding components of the T-cell receptor, and known cytotoxic mediators (Davoli et al (2017), see Supplementary Table 4) in the same TIGER networks. A striking finding was that *CCL19* is significantly negatively associated with LOY (Pearson = *rho* = − 0.28; FDR-corrected *p* = 0.0038); *CCL19* can chemoattract T- and B-cells (Kim et al, 1998) and has been shown to mediate potent immune responses against tumors in mouse models of lung cancer (Hillinger et al, 2006). This suggests a potential causal link between LOY and decreased immune infiltration in tumors. These regulatory network analyses further clarify the mechanism by which LOY may be selected for–that LOY may alter the regulation of CTAs and immune-response-related genes, altering the infiltration of white blood cells in the tumor and aiding tumor cell immune evasion.

#### Gene regulatory sub-networks activated with LOY reflect influence on tumor progression

Although a node-based analysis of gene regulatory network models can identify elements associated with phenotypic features, it does not fully capture the patterns of interconnectivity represented in the network model. “Active” sub-network analysis integrates gene expression data with the network structure to pinpoint areas where there is a notable deviation in activity compared to a control or baseline condition. Following the approach of Dittrich et al (2008) (see Methods 2.4), we identifed active sub-networks in the TIGER LUAD GRN that are associated with LOY. We found 337 nodes with positive scores resulting in a bipartite active sub-network consisting of 210 nodes (10 transcription factors, 200 genes) and 233 TF-gene edges. The key network attributes of the ten TFs in the active sub-network are summarized in Supplementary Table 5.

The highest degree TF was NR3C2 (weighted absolute outdegree: 23.96), followed by KLF15 (16.85), for which the bipartite sub-network is shown in Figure 4b. Both NR3C2 and KLF15 have a strong negative association with LOY (blue node coloring), indicating lower expression among individuals who have lost the Y chromosome. KLF15 has been shown to regulate pathways that intersect with immune responses, potentially contributing to immune evasion mechanisms by altering the tumor microenvironment and immune cell infiltration (Shu et al, 2024). NR3C2 is known primarily for its role in regulating electrolyte balance but also has been found to play a part in cancer biology, including immune evasion (Sun et al, 2024b). In lung adenocarcinoma, NR3C2 can influence immune response by affecting the expression of immune-regulatory genes. This modulation can help the tumor evade immune surveillance by altering the activation and infiltration of immune cells (Chen et al, 2023; Shu et al, 2024). In most cases, their targets also have a negative association with LOY, consistent with the network trend for NR3C2 and KLF15 as activators (primarily red colored edges in Figure 4b). The active subnetwork also contains the TF GATA3, which also decreased in expression with LOY (effect= −0.34, *p* = 1.52 ∗ 10^−6^, FDR=1.19 ∗ 10^−4^). GATA3 plays a critical role in the development and function of T cells and other cell types, including playing a role in cell differentiation and proliferation (Wan, 2014). Aberrant expression of GATA3 can alter the regulation of cytokines and other signaling molecules that modulate the immune system’s ability to recognize and attack tumor cells (Anichini et al, 2020).

Only two TFs, ZC3H8 and ADNP, increased in expression with LOY. The association between ADNP and LOY is not statistically significant (effect = 0.03, p = 0.63) and the heuristic algorithm likely selected ADNP for inclusion in the active subnetwork because of the strong associations of several of its target genes with LOY. However, high levels of ADNP expression have been linked with poor prognosis and enhanced tumor progression, and it is thought to play some role in immune modulation (Wang et al, 2023). ZC3H8 exhibits a significant positive association with LOY (effect = 0.39, *p* = 1.91 ∗ 10^−7^, FDR=6.18 ∗ 10^−5^) and serves as a repressor of six of the seven of its target genes in the active GRN subnetwork (Figure 4b). ZC3H8 is a known repressor of *GATA3* (Schmidt et al, 2018), consistent with the relationship we see between their expression. Thus, ZC3H8 likely influences immune regulation as expected; its increased expression has been linked to poor outcomes in LUAD (Sun et al, 2024a).

Closer investigation of the active subnetworks of the TFs KLF15 and NR3C2 further supports the hypothesis of LOY-related change in regulatory control affecting the immune environment. *SUSD2* showed one of the highest positive regulatory controls by NR3C2, while having a negative association with LOY. SUSD2 suppresses CD8+ T cell antitumor immunity by targeting IL-2 receptor signaling (Zhao et al, 2022). *CENPN* and *CDC20* showed the highest negative regulation in the NR3C2 active sub-network, while both being postively correlated with LOY. *CENPN* was reported to be associated with genetic markers of immunomodulators in various cancers and is differentially expressed across molecular and immune sub-types (Zhang et al, 2025; Jing et al, 2025). CDC20 was reported to regulate apoptosis, immune microenvironment, and tumor angiogenesis (Xian et al, 2023).

We also investigated the properties of the 200 targeted genes in the active sub-network using an over-representation analysis. We used the Canonical Pathways (CP) subset of the MolSigDB gene set repository as our annotation source; this set includes the KEGG-Medicus, REACTOME, WikiPathways, and KEGG legacy gene sets. The background was set to all genes in the full TIGER network. Cytochrome p450 (CYP) pathways were consistently found to be significant when using all three annotation sources: the KEGG “drug metabolism cytochrome p450” pathway (FDR = 0.013), the WikiPathways “oxidation by cytochrome p450” pathway (FDR = 0.034), and the REACTOME “cytochrome p450 arranged by substrate type” pathway (FDR = 0.035). Other top pathways enriched among the targeted genes, consistent with the activation of CYP genes, included the REACTOME “phase 1 functionalization of compounds” (FDR = 8.6 ∗ 10^−5^) and “biological oxidations” (FDR = 0.010) and the KEGG “fatty acid metabolism” pathway (0.014). Full results of the over-representation analysis, including lists of overlap genes in each pathway, are available in Supplementary Table 6.

CYP enzymes play significant roles in the metabolism of both endogenous and exogenous compounds, influencing not only cancer progression and treatment outcomes but also immune responses within the tumor microenvironment. CYP enzymes are known to play significant roles in xenobiotic and drug metabolism, and the regulation of CYP enzymes by pro-inflammatory cytokines has been implicated in cancer prognosis and therapeutic response (Stipp and Acco, 2021). In lung adenocarcinoma, sex differences in CYP expression have been associated with the increased risk of developing lung adenocarcinoma in females (Oyama et al, 2007).

### LOY and immunotherapy

The LOY-driven changes we observed in tumor cell gene expression and TIME composition led us to speculate that LOY may affect response to treatment with cancer immunotherapy agents. We explored the association between LOY and response in a dataset of 832 individuals with LUAD (kindly provided by Caris Life Sciences), consisting of bulk gene expression data for cohorts treated with pembrolizumab (pembro, N=832), with response assessed through survival after treatment. Further available data for cohorts treated with nivolumab (N=125) and atezolizumab (N=55) were not considered for in-depth analysis due to low statistical power caused by an order of magnitude smaller sample size. We provide hazard ratio plots based on LOY and CTA expression across all treatments in Supplementary Figure 5. In compliance with HIPAA, no individual-level information was provided so the analyses presented here use summary statistics for data partitions such as high vs low LOY (see Methods).

Because the Caris data did not have individual-level Y chromosome information, we could not calculate LOY directly; instead used the expression of *RPS4Y1* as a proxy. We chose *RPS4Y1* because it was the Y-chromosome gene we found to have the greatest weight in pc-LOY and had the highest single-gene correlation with LOY in bulk sequencing data (Pearson *ρ* = 0.84 in male individuals in TCGA-LUAD). Consequently, we used the median *RPS4Y1* expression to define high and low LOY groups. We compared estimated TIME composition in those groups using quanTIseq; consistent with our analyses in other datasets, we found significant differences between high and low LOY for M1 and M2 macrophages, neutrophils, monocytes, B-cells, and CD8+ T-cells (FDR-adjusted p-value *<* 0.05; Supplementary Table 7). In the Caris data, we found that M1 macrophages increase with LOY; previous work has shown M1 macrophage infiltration in tumors to be predictive of immunotherapy response in NSCLC (Wu et al, 2024).

We compared treatment outcomes for individuals with and without LOY. For pembro-treated individuals, LOY was associated with better outcomes as measured by the hazard ratio between the two partitions (HR = 0.831, 95% CI = 0.722-0.956, *p* = 0.0095), which is consistent with Abdel-Hafiz et al (2023), which reported improved response to PD-1 antibodies in a mouse model of LOY.

Because we had seen that LOY was correlated with CTA expression and that the expression of 26 CTAs was correlated with survival in TCGA, we wanted to determine whether the expression of these CTAs was predictive of immunotherapy response. Data were available from Caris for 25 of the predictive CTAs, and for each, we partitioned each treatment group into low- and high-expression for that CTA according to cutpoints at the 75th percentile. The relationship between CTA expression and survival was generally treatment-specific, with four of the 25 measured CTAs showing a significant association with survival in the pembro treatment group (*p <* 0.05; Figure 4c). These findings provide evidence that LOY can affect immunotherapy outcomes, likely by modifying the TIME through CT antigen expression. Hence, an individual’s likely treatment response could be estimated using the amount of LOY or expression of specific CTAs, and treatment adjusted to increase the likelihood of a positive outcome.

## 4 Discussion

Despite substantial evidence that there are clinical differences between biological males and females in disease risk, development, and response to therapy, the role of allosomes and the molecular processes associated with them have only recently begun to be studied in earnest in the context of lung cancer (Lopes-Ramos et al, 2018, 2020; Arnold and Disteche, 2018). The role that the male Y chromosome plays in cancer risk, development, and response to therapy was also poorly understood despite growing evidence that LOY in solid tumors is associated with poor outcomes and that LOY may be a cancer driver (Brown and Machiela, 2020; Cáceres et al, 2020; Qi et al, 2023; Müller et al, 2023).

Our analyses provide support for the hypothesis that LOY is not just a passive driver but that it is selected for lung adenocarcinoma development in biological males. We first demonstrated that LOY is associated with alterations in patterns of expression of Y-chromosomal genes. While not surprising, this observation provides a means of inferring LOY in datasets that do not include direct sequencing of the Y chromosome. Armed with this information, we tested whether the LOY we observed was due to chromosomal loss in tumor cells or infiltrating lymphocytes. We found that the LOY in tumors is confined to cancer cells, not the lymphocytes present in the tumor. This analysis was critical because it is known that LOY occurs in the white blood cells of aging males, and so the LOY in the tumor could have been confounded by LOY in the infiltrating immune cells. Next, we examined the composition of white blood cells in tumors as a function of Y chromosome loss. We discovered that there were associated changes in immune cell composition, including increased levels of M1 macrophages that are well correlated with LOY. We also tested for expression of cancer/testes antigens (CTAs) and found that CTA expression was correlated with both LOY and changes in the tumor immune microenvironment (TIME).

Collectively, this set of facts suggests a simple model of selection for LOY in solid tumors in biological males. As cancer develops, immune cells recognize CTAs on tumor cells and attach to them. However, some tumor cells lose their Y chromosome, resulting in diminished expression of CTAs. This allows them to avoid immune attack, selecting for those cells that have lost the Y chromosome. As tumor cells that have experienced LOY begin to dominate the tumor cell population, the overall composition of the TIME starts to change, resulting in the process of tumor evolution that is selectively enriching for LOY. This model is, in fact, consistent with the report that LOY in bladder cancer is a driver of tumor growth and is accompanied by a change in the tumor immune environment that facilitates immune evasion (Abdel-Hafiz et al, 2023; Chen et al, 2025).

Our proposed model not only sheds light on the role of the LOY in LUAD through its role in immune invasion but also suggests that LOY or CTA expression patterns may be relevant to understanding immunotherapy response. Indeed, CTA expression has been reported to have a strong association with survival in LOY-affected individuals; some have suggested using CTA expression as a biomarker for immunotherapy (Gjerstorff et al, 2015; Al-Khadairi and Decock, 2019; Meng et al, 2021; Seager et al, 2024), and CTAs themselves have been suggested as targets for immunotherapy (Saito et al, 2016; Simister et al, 2022). Although we were only able to draw firm conclusions for pembro, the results for nivo and atezo suggest further investigation of whether LOY can be used to determine optimal immunotherapy treatment.

A consistent observation across our studied datasets, including immunotherapy data, is that M1 macrophages increase with LOY. M1 macrophages have been reported to be associated with better immunotherapy treatment response in lung cancer (Wu et al, 2024), which may help explain why the changes in the TIME associated with LOY result in better treatment effects. There is also evidence that the presence of M1 macrophages improves treatment efficacy for multiple immunotherapeutic treatments (Duan and Luo, 2021; Huo et al, 2023). As such, LOY could also serve as a biomarker for selecting therapies with responses linked to the presence of M1 macrophages. Within the studied immunotherapy data, we further find that *XAGE1B* is a good marker for pembro treatement in LUAD showing exceptionally high hazard ratio. Immunotherapy treatments explicitly targeting CTAs such as *XAGE1B* (Saito et al, 2016) could further be considered as new therapeutic targets, particularly if those treatments are conditioned on whether a subject shows LOY.

What we have learned about the role that LOY plays in lung tumor development and in affecting changes in the immune environment provides a strong motivation for a more complete study of the role that allosomes play in disease processes. The allosomes have long been neglected in the study of health and disease (Sun et al, 2023). While LOY is gaining recognition as a marker of certain cancers and male aging, its functional role is still relatively unknown. Our findings point to a compelling but straightforward model for why LOY is prevalent in male solid tumors: selective pressure. As tumor cells lose their Y chromosome, they alter the expression of CTAs and the composition of the TIME. These changes allow cells that have undergone LOY to better evade immune surveillance, giving them an advantage over tumor cells that retain the Y chromosome. This also strongly suggests that an exploration of the mosaic loss of X could further our understanding of sex differences in immune response (Klein and Flanagan, 2016) and may shed light on sex-biased processes that are active in the tumors of biological females.

As is true of all studies, this study is limited by the quantities and types of data that are available or that we can practically generate. Lacking that data, we are confident in the model because of the large number of lines of evidence derived from multiple sources that support it. Cancer development and progression is fundamentally a process driven by selection in which tumor cells gain an advantage over normal cells and expand in numbers. As put quite eloquently by Martincorena et al (2017), “By definition, cancer genes are genes under positive selection in tumor cells.” It is simply logical to extend this concept to subpopulations of tumor cells defined by their genetic makeup—including whether or not they possess a Y chromosome.

Similarly, the conclusions we reached regarding the association between LOY and immunotherapy response are based on a relatively small number of individuals with LUAD who received immunotherapy. Recall that for two of the drugs, nivo and atezo, there was insufficient data to draw significant conclusions. While the results for pembro were statistically significant, they suggest further independent validation in larger independent datasets. Nonetheless, the model here is also compelling—LOY clearly influences CTA expression, tumor progression, immune processes, and the TIME, and M1 cell infiltration—providing a strong argument for the potential use of LOY as a marker for immunotherapy response in males with lung adenocarcinoma.

## Acknowledgments

This work was supported by grants from the National Institutes of Health: JF, KHS, CC, VF, ES, PM, MBG, CMLR and JQ were supported by R35CA220523; MBG and JQ were also supported by U24CA231846; JQ received additional support from P50CA127003; JQ and DLD were supported by R01HG011393; KHS and DLD were supported by P01HL114501; KHS was supported by T32HL007427; DLD was supported by K24HL171900; CMLR was supported by K01HL166376; CMLR and ES were also supported by the American Lung Association grant LCD-821824.

The results published here are in part based upon data generated by the TCGA Research Network: https://www.cancer.gov/tcga. The Genotype-Tissue Expression (GTEx) Project was supported by the Common Fund of the Office of the Director of the National Institutes of Health, and by NCI, NHGRI, NHLBI, NIDA, NIMH, and NINDS. Immunotherapy data were provided in aggregate by Caris Life Sciences; JQ serves on the Caris Scientific Advisory Board but was not compensated in any way for the analysis presented here.

## Data Availability Statement

We provide code and documentation for generation of the presented analysis and anonymized and aggregated information on immunotherapy results specifically used in our study at https://github.com/QuackenbushLab/LOY_lung_cancer. Other data is publicly available, we provide all data sources along with identifier in the Methods sections.

