## Supplementary Figure for "Selective loss of Y chromosomes in lung adenocarcinoma modulates the tumor immune environment through cancer/testis antigens"

November 4, 2025

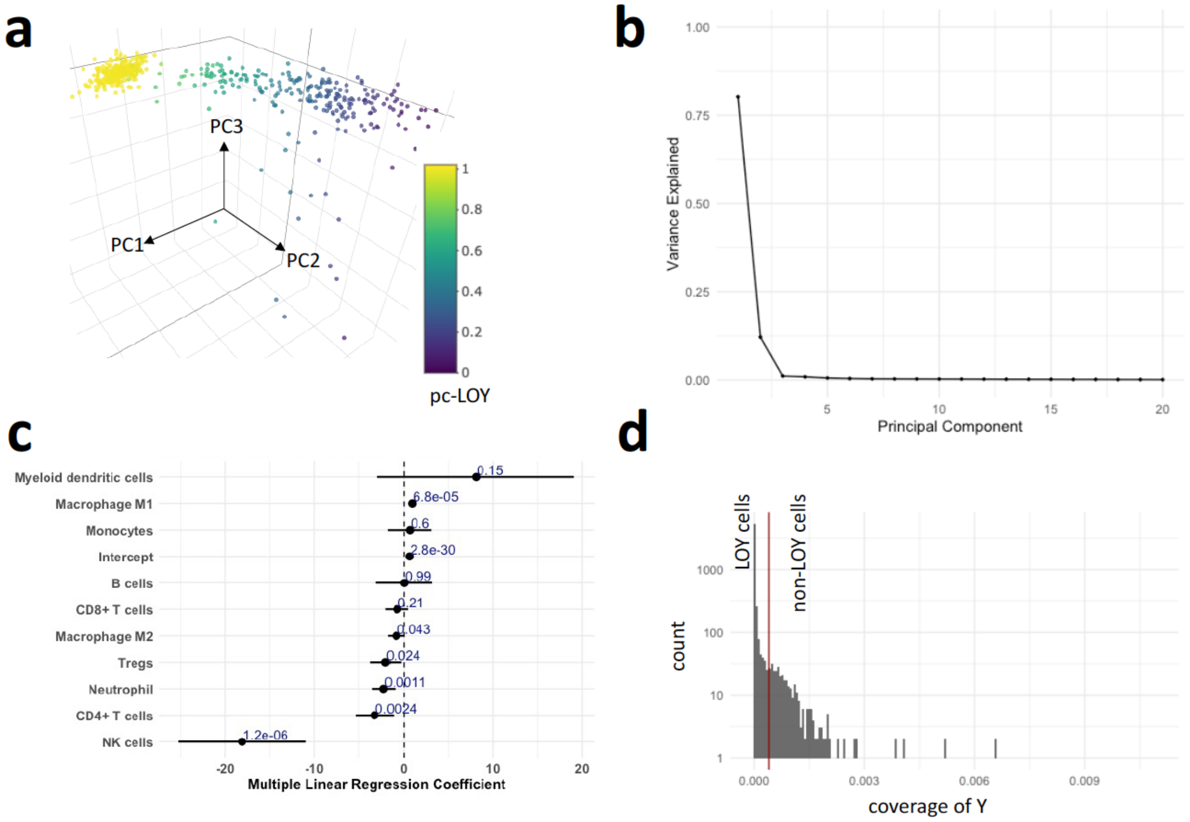


Supplementary Figure 1: *Additional analysis on LOY.* **a** The first three principal components of TCGA-LUAD, which are used for pc-LOY construction. Coloring by resulting pc-LOY, the cluster on the top-left are the female reference samples. **b** Scree plot of first 20 principal components on TCGA-LUAD. **c** Forest plot for coefficients of Quantiseq immune cell type fractions in a least squares LOY fit. Visualized are coefficient estimates with standard error and p-value. **d** Coverage distribution of the Y chromosome across single epithelial cells in HTAN-LUAD. LOY threshold $(4 * 10^{-4})$ in red.

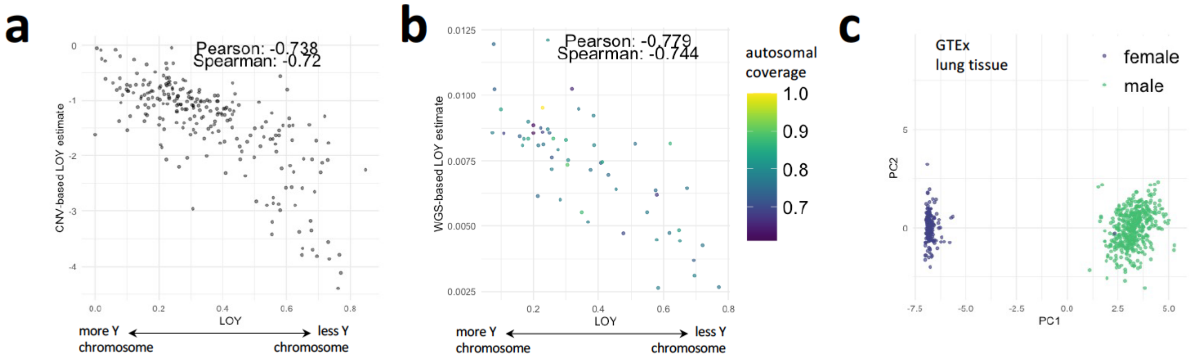


Supplementary Figure 2: *Different measures of LOY and covariates.* **a** TCGA LUAD bulk gene expression-based LOY estimate (x-axis) against copy number variation (CNV) based LOY estimate (y-axis). Average CNV across all Y segments from TCGA LUAD data for which matching gene expression exists. **b** TCGA LUAD bulk gene expression-based LOY estimate (x-axis) against whole genome sequencing (WGS) based LOY estimate (y-axis). Coverage of the Y chromosome relative to overall genome coverage in WGS data, coloring corresponds to autosomal coverage, normalized by maximum observed autosomal coverage across all samples. **c** First two principal components of Y gene expression in GTEx lung tissue.

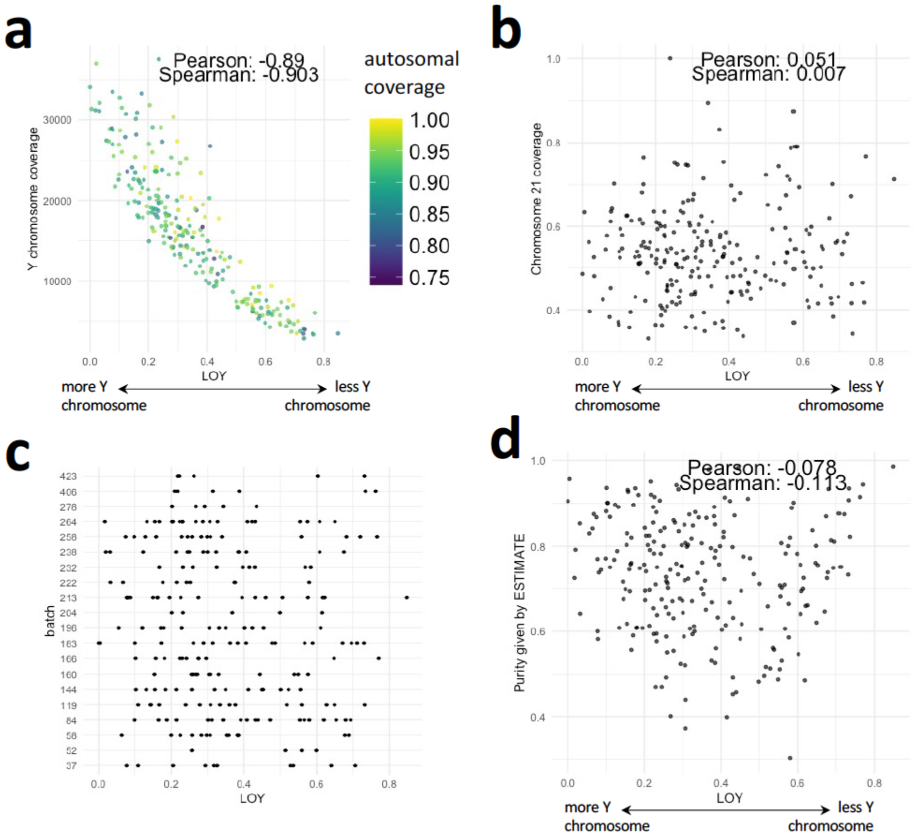


Supplementary Figure 3: *Bulk LOY coverage analysis and single-cell estimates.* **a** Overall Y chromosome coverage in TCGA LUAD bulk sequencing data (y-axis) against the LOY estimate (x-axis), colored by autosomal coverage in bulk sequencing. **b** Coverage of chromosome 21, a small and gene-deserted chromosome, relative to the LOY estimate in TCGA LUAD. **c** Reported batch ID against LOY estimate in TCGA LUAD. **d** Sample purity estimate through ESTIMATE method against LOY estimate in TCGA LUAD.

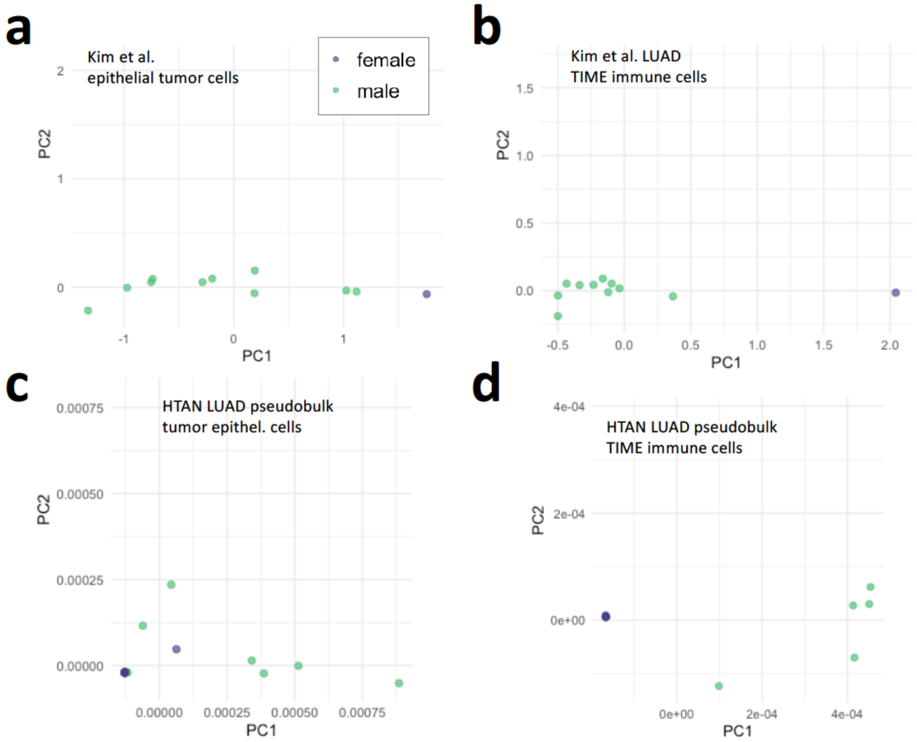


Supplementary Figure 4: *Gene set enrichment in LOY cells.* textbfa-b First two principal components on Y genes of pseudobulked gene expression of annotated epithelial cells (left) and immune cells (right) for LUAD data of Kim et al. **c-d** First two principal components on Y genes of pseudobulked gene expression of epithelial cells (left) and immune cells (right) for LUAD data from the HTAN.

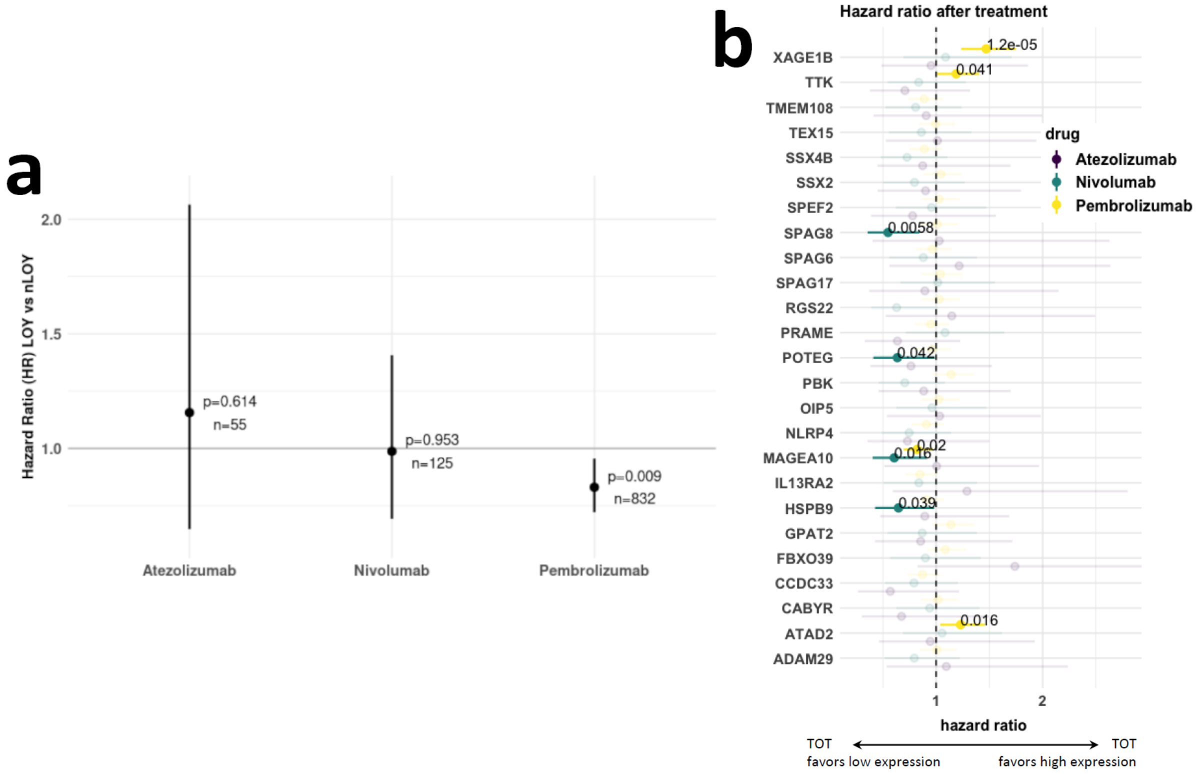


Supplementary Figure 5: *Immunotherapy response in the context of LOY for different treatment cohorts.* **a** Hazard ratio (y-axis) comparing the group with LOY compared to that showing no LOY group; groups were assigned using an RPS4Y1-based LOY estimate and HR measures as high expression versus low expression of RPS4Y1. **b** Hazard ratio (x-axis) for male LUAD cohorts treated with specific drug (color) based on median expression value of CTA (y-axis, low expression group vs high expression group). HR values with confidence intervals and p-values in bold, all insignificant observed HRs are transparent for clarity of the results.
